# Universal rapid RNA-based quantification of toxigenic *Alexandrium* species (Dinophyceae) using quantitative recombinase polymerase amplification

**DOI:** 10.64898/2025.12.28.696751

**Authors:** Ioannis Markopoulos, Isavella Papadopoulou, Christos Chantzaras, Katherine Hartle-Mougiou, Frédéric Verret, Electra Gizeli, Martha Valiadi

## Abstract

Harmful algal blooms caused by toxigenic *Alexandrium* species pose recurrent risks to coastal ecosystems and public health, yet current monitoring approaches rely on microscopy and laboratory-based toxin analysis with limited capacity for rapid, functional early warning. Here, we present a universal quantitative reverse-transcriptase recombinase polymerase amplification (qRT-RPA) assay targeting the *sxt*A4 transcript, an essential gene in saxitoxin biosynthesis. The RPA chemistry is isothermal with a low running temperature, making it suitable for portable, on-site testing. The assay was designed against a conserved *sxt*A4 region and validated using isolated *sxt*A4 amplicons from multiple *Alexandrium* species, synthetic DNA and RNA templates, and total RNA of *Alexandrium minutum* as a widespread reference species. The assay achieved uniform amplification kinetics across species, a limit of detection below 10³ synthetic RNA copies and 0.1 ng total RNA per reaction for *A. minutum*, and a runtime of less than 15 minutes. The assay selectively detected *sxt*A4 transcripts from toxigenic *Alexandrium* strains and showed no cross-reactivity with non-target phytoplankton. A limit of detection that is relevant to early warning for *Alexandrium* species (under 20 cells per reaction) was retained in complex RNA matrices. Mock samples prepared by spiking cultured cells into natural seawater also demonstrated detection at field-relevant concentrations. These results establish qRT-RPA as a rapid, RNA-based, functionally informative molecular tool that provides a foundation for portable, early-warning monitoring of potentially toxigenic *Alexandrium* blooms.

## 1 Introduction

Harmful Algal Blooms (HABs) are posing increasing ecological and socioeconomic pressure in coastal areas worldwide. Their global expansion is attributed to anthropogenic nutrient enrichment and ocean warming (Dai et al., 2023; Gobler et al., 2017; Griffith et al., 2019; Hayashida et al., 2020; Rabalais et al., 2007; Yu et al., 2023). Warming temperatures can promote bloom initiation (Anderson et al., 2021a) accelerated cell growth rates and prolonged bloom seasons (Anderson et al., 2021a; Gobler et al., 2017; Griffith et al., 2019) especially in temperate regions (Dai et al., 2023), whilst also allowing their intrusion into previously unaffected polar areas (Anderson et al., 2021a). The consequences of HABs range from ecosystem-scale deoxygenation to the accumulation of potent toxins that threaten food security and human health (Anderson et al., 2021a; Anderson et al., 2021b). Despite progress in ocean observing systems, early-warning capacity often remains limited, particularly in regions where toxic events develop rapidly or surveillance infrastructure is sparse.

The dinoflagellate genus *Alexandrium* is responsible for HABs worldwide. Species of this genus produce an array of toxins as secondary metabolites, being most well-known for the production of saxitoxins. These cause Paralytic Shellfish Poisoning (PSP) in humans upon consumption of contaminated shellfish ((Anderson et al., 2012; Gribble et al., 2025; Lewis et al., 2018; Suleiman et al., 2017). Most regulatory monitoring programs still rely on a dual structure: (i) microscopic identification and enumeration of cells in water samples, and (ii) chemical toxin analysis in shellfish tissue (Parks et al., 2020; Turner et al., 2014; union, 2004). The first assesses putatively toxic cell abundance, while the second measures saxitoxin and congeners which are converted to toxicity equivalence factors (TEFs) (Commission Implementing Regulation (EU) 2021/1709, (Commission., 2021). Concerned countries like the UK follow this regime with practical thresholds of 400 STX eq. (Turner et al., 2014) in shellfish flesh or 40 - 100 *Alexandrium* cells L^-1^ of water triggering intensified monitoring frequency ((Parks et al., 2020; Senovilla-Herrero et al., 2025). In other locations the thresholds may be much higher if cells are less toxic; there is considerable variability in toxicity level regionally, interspecifically and intraspecifically (Geffroy et al., 2021; Sung et al., 2025). While these methods underpin long-established monitoring frameworks, they present significant operational constraints. Microscopy cannot reliably distinguish between toxic and non-toxic strains of morphologically similar taxa and provides no functional information on active toxin production at the time. Meanwhile, toxin chemical analyses by HPLC (Standard EN 14526) requires centralized laboratories, limiting sampling frequency and increasing turnaround time to minimum of 3 days. Together, these limitations hinder early warning and intervention, increasing the likelihood of food contamination and ecological damage.

Molecular amplification-based methods represent an alternative to microscopy analysis, offering higher sensitivity, faster time-to-result, species-level resolution and quantification of specific cellular functions (John et al., 2007; Kon et al., 2015; Murray et al., 2019). A wealth of DNA-based qPCR assays have been developed to enumerate specific toxic species (Dai et al., 2020; Erdner et al., 2010; Gao et al., 2015a; Gao et al., 2015b; Garneau et al., 2011; Hatfield et al., 2019; Ruvindy et al., 2018; Toebe et al., 2013; Vandersea et al., 2017). Most existing assays target the small or large ribosomal genes of specific *Alexandrium* species to estimate abundance, whereas several qPCR assays targeting genes of the saxitoxin cluster enable direct assessment of the population’s toxigenic potential (Gao et al., 2015b; Han-Sol et al., 2024; Murray et al., 2019; Murray et al., 2011; Penna et al., 2015; Savela et al., 2016; Stüken et al., 2013). Despite these advances, the transition from traditional light microscopy to molecular monitoring for HAB surveillance has been slow to materialize. While qPCR is the gold standard of molecular detection due to its cost-effectiveness and high throughput, it is mainly applied in a laboratory setting due to the requirement for high-power consuming thermal cycling.

HAB surveillance will significantly benefit from on-site or *in situ* monitoring offering real-time detection. In contrast to qPCR, isothermal amplification can achieve molecular quantification in remote field settings, using portable battery-powered analyzers (e.g. Pebble R by BIOPIX-T; TwistAmp by TwistDx). In particular, recombinase polymerase amplification (RPA) operates at a low, stable temperature (37 – 42 °C), has high tolerance to impurities and a fast time-to-result in less than 15 minutes (Tan et al., 2022). RPA chemistry can be used for qualitative detection using lateral flow strips (Liu et al., 2023; Yao et al., 2024; Yu et al., 2025) and quantitative analysis by incorporating fluorescent probes into the reaction (Tan et al., 2022). RNA can be amplified in one step simply by adding reverse transcriptase in the RPA reaction. These features make RPA a versatile and fast amplification technology that is suitable for portability, and can be applied to quantify functional genes in field-monitoring applications.

Saxitoxin biosynthesis in dinoflagellates is linked to a family of genes homologous to the cyanobacterial *sxt* cluster (Kellmann et al., 2008; Stüken et al., 2011). The *sx*tA4 domain is recognized as essential for toxin production and its expression is strongly correlated with intracellular toxin content (Geffroy et al., 2021; Murray et al., 2015). This makes the *sxt*A4 gene domain an appropriate target for quantifying toxic populations rather than simply detecting genus presence.

In this study, we use the HAB forming, saxitoxin-producing dinoflagellate genus *Alexandrium* as a case-study, to demonstrate the utility of qRT-RPA for RNA-based quantification of metabolically active toxigenic cells. We present optimization strategies for qRT-RPA to achieve an equivalent performance across four *Alexandrium* species, despite minor sequence variation. We the use *Alexandrium minutum* as a reference species to benchmark limits of detection and standard curves for sample quantification. We demonstrate the assay performance in realistic conditions using simulated samples. To our knowledge, this is the first report of an RNA-based quantitative RPA assay for a harmful algal bloom species. We therefore provide a methodological foundation for transitioning HAB molecular detection into portable-ready, rapid-response monitoring, positioning RPA as a robust approach for HAB surveillance and early warning.

## 2 Materials and Methods

### 2.1 Culture maintenance and growth experiments

Culture isolates of dinoflagellates and other phytoplankton species were obtained from the Roscoff Culture Collection (RCC, France), the Culture Collection of Algae and Protozoa (CCAP, UK), and the Culture Collection of Harmful and Environmental Marine Microalgae (CCVIEO, Spain). Strains were maintained in L1 culturing media (CCAP, UK) in polypropylene flasks (Starstedt), at 18°C, with a light : dark cycle of 16 : 8 hours and an irradiance of approximately 100 μmol m^−2^ s^−1^. Cells were kept in exponential growth phase by frequent subculturing and monitoring the cell concentration using a Sedgewick-Rafter counting chamber and a brightfield microscope (Olympus CX-43).

### 2.2 Nucleic acid isolation

Cell material was collected for nucleic acid extraction by either pelleting by centrifugation at 4000g for 10 mins, or by vacuum filtration onto a 3.0 μm pore size polycarbonate membrane (Nuclepore, Whatman, USA). Samples were flash frozen in liquid nitrogen and stored at −80 C until further processing. Immediately prior to nucleic acid isolation, cells were disrupted using liquid nitrogen and grinding with a pestle. DNA was extracted using the DNeasy Plant Mini Kit (Qiagen, Netherlands). RNA was extracted using the GRS Total RNA kit – plant (GRiSP, Portugal), residual DNA was removed from RNA using Turbo DNAse treatment (Thermo Fisher Scientific, USA) and re-purified using the Monarch RNA cleanup kit (NEB, USA). DNA and RNA purity was assessed spectrophotometrically (Nanodrop One, Thermo Fisher Scientific, USA) and quantity was measured using a Qubit 4 instrument (Thermo Fisher Scientific, USA). Additionally, RNA integrity was also assessed using a Bioanalyzer 2100 with Plant RNA pico reagents (Agilent, USA) electrophoresis.

### 2.3 RPA primer and probe design

*Sxt*A4 gene sequences from all dinoflagellate species were retrieved from GenBank and aligned through NCBI Blastn. A consensus sequence was generated for each species using EMBOSS cons (https://www.ebi.ac.uk/jdispatcher/msa/emboss_cons). The most conserved regions for assay design covering all *Alexandrium* species were identified manually.

RPA primers (Eurofins, Germany and IDT, USA) were designed according to TwistDx’s RPA assay design manual (GC content: 20–70%, length: 30–35 nt, Tm: 50–100 °C), using Primer-BLAST on *Alexandrium minutum sxt*A4 gene (GenBank accession number KM438016.1). Primer candidates were evaluated for 1) potential self- and cross-dimer formation using the Multiple Primer Analyzer (Thermo Fisher Scientific), 2) stable secondary structures using Mfold (unafold.org) and 3) cross-reactivity using Blastn. Where potential primer dimers were detected at the 5’ or 3’ end, the primer sequence was truncated by 1-2 nucleotides.

The RPA probe was designed according to TwistDx’s RPA assay design manual and produced by Biomers (Germany). The design criteria were expanded based on the literature to specifically minimize the G-content of the probe and reduce the distance between fluorophore and quencher to a minimum (Liu et al., 2019). A custom script identified regions of 47-nucleotides on both the forward and reverse strands of the target sequence, ensuring a thymine was at position 31 as recommended by the manufacturer. Probes predicted by the Multiple Primer Analyzer to form dimers on the abasic site were excluded to prevent non-specific fluorescence emission. The final selected primers and probe are shown in **Table 1**.

**Table 1.**
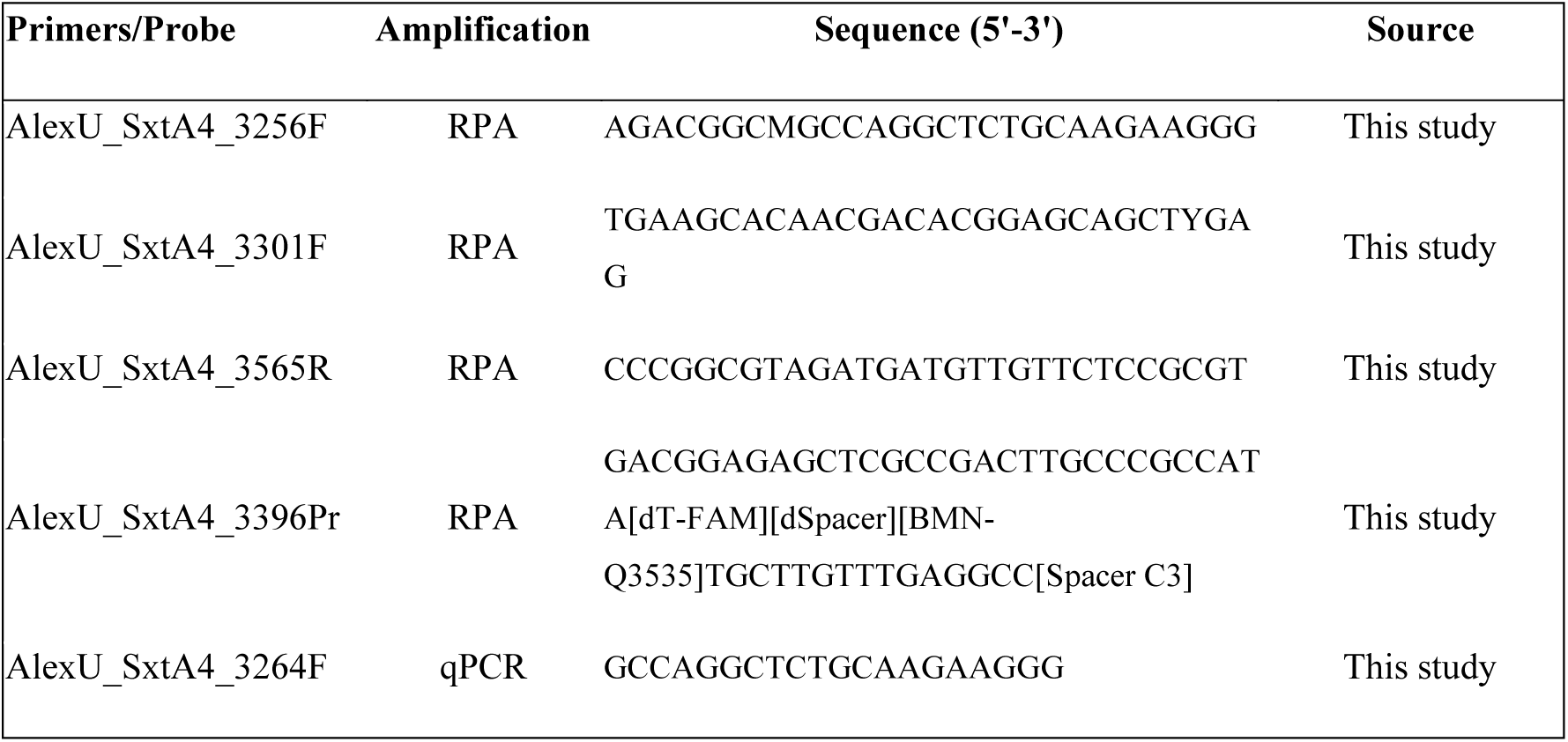
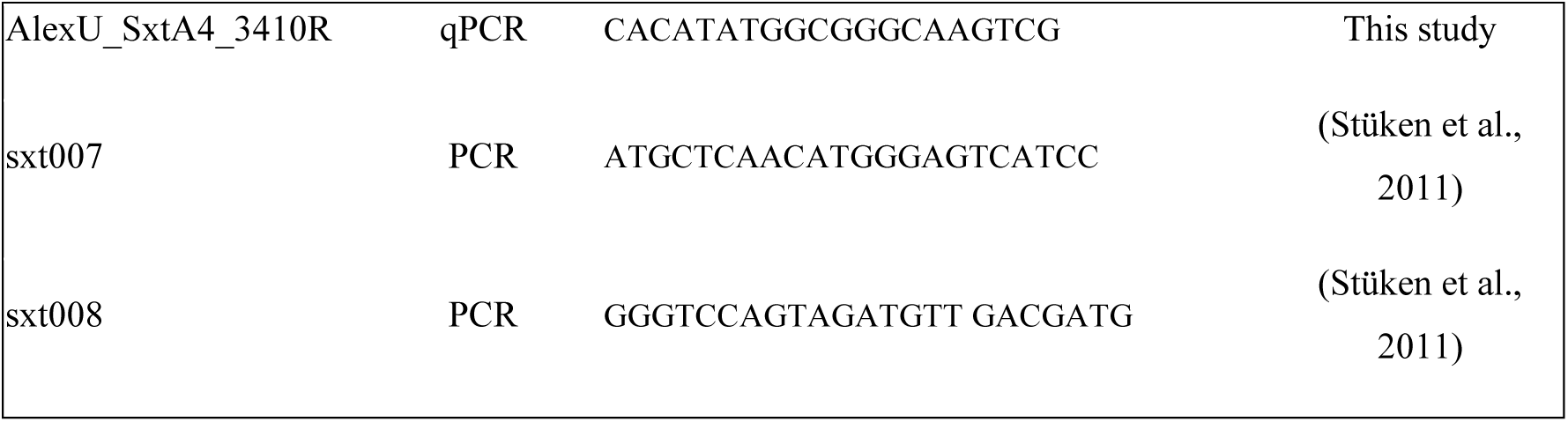
Primer and probe sequences used in this study.

### 2.4 PCR amplification of *Sxt*A4 from gDNA of cultured isolates

To confirm the presence of the *sxt*A4 gene in *Alexandrium* cultured strains VGO0650, VGO0657, CCAP 1119/48 and FE126, we used a touch-up PCR protocol utilizing *sxt*A4 specific primers (Stüken et al., 2011) (**Table 1**). All PCRs were run at 25 μL reaction volumes containing 1x GoTaq G2 green Master Mix (Promega, USA), 5% v/v DMSO, 0.4 μM Forward and Reverse primers. PCR cycling was carried out as follows: 95°C for 3 minutes, 5 x (95°C for 30s, 58°C for 45s, and 72°C for 50s), 5 x (95°C for 30s, 60°C for 45s, and 72°C for 50s), 27 x (95°C for 30s, 62°C for 45s, and 72°C for 50s), 72°C for 5 minutes. Results were visualized by electrophoresis on a 1.5% agarose gel.

Where *sxt*A4 was detected, we repeated the PCR in triplicate with a high-fidelity polymerase to recover the amplicons for sequencing in order to verify their identity against the reference sequence of the strain on NCBI. All PCRs were run in 25 μL reaction volumes containing 1x KAPA HiFi HotStart ReadyMix (Roche, Switzerland), 5% v/v DMSO, 0.4 μM Forward and Reverse primer and water adjusted for the volume of the template. The PCR cycle was: 95°C for 3 minutes, 35 x (95°C for 20s, 60-65°C for 15s, and 72°C for 50s), 72°C for 2 minutes. After gel electrophoresis the bands were excised, purified using the NucleoSpin Gel and PCR Clean-up kit (Macherey-Nagel, Germany), both strands were sequenced by Genewiz (Azenta Life Sciences, USA).

### 2.5 Recombinase Polymerase Amplification

Quantitative RPA (qRPA) and Reverse Transcriptase-RPA (qRT-RPA) reactions were performed following the manufacturer’s instructions for the TwistAmp Exo kit (TwistDx, Cambridge, UK). Real-time, exonuclease-assisted RPA reactions were run at 40°C, at a final volume of 25 μL per reaction and detected on a CFX Opus 96 Real-Time qPCR system (BioRad, USA) at 15 second intervals. Initially, RPA reactions were setup following the manufacturer’s instructions and further optimized as shown in section 3.2. Briefly, for a 50 μL reaction which was then split in half, a rehydration master mix was prepared containing 29.5 μL rehydration buffer, variable concentrations of forward primer (240 – 600 nM), variable concentrations of reverse primer (240 – 600 nM), variable concentrations of probe (90 – 150 nM), 1 μL Superscript IV for RNA templates (Thermo Fisher Scientific, USA) and water up to the required volume. The freeze-dried enzyme pellet was reconstituted with rehydration master mix and split into two individual 25 μL reactions, followed by template addition. MgOAc at variable concentrations (14 – 30 mM) were added to the flat cap of each reaction. The introduction of MgOAc into each reaction and the subsequent mixing were performed by simultaneously spinning down, vortexing and spinning down again all reactions together, before being placed in the qPCR instrument. Measurements of Time to Positivity (TTP) were calculated as the timepoint at which the fluorescent signal crossed the fluorescence threshold, using OriginPro’s (2024) Intersect Gadget. The threshold for each reaction was set as the mean fluorescence signal plus three times the standard deviation of all the negative (no template) control measurements.

### 2.6 Assay performance across multiple *Alexandrium* species

Firstly, the assay’s performance was tested across four species *A. tamarense* (RCC292)*, A. ostenfeldii* (CCAP 1119/53)*, A. pacificum* (CCAP 1119/52)*, A. minutum* (VGO0657). For this test, we used *sxt*A4 amplicons generated from the cultured cells and amplified by qRPA (omitting reverse transcriptase). This step reveals any difference in amplification based on the target sequence, without requiring costly production of synthetic RNA for each representative species, with minor sequence differences. The amplicon concentrations were determined spectofluorometrically (Qubit, Invitrogen, USA) and serially diluted for testing over 4 orders of magnitude. A synthetic DNA fragment of the target region representing *A. minutum* consensus sequence, plus 20 bp flanking either side (IDT, USA) was also included for additional quantification accuracy.

### 2.7 Assay sensitivity, quantitative range and specificity

The assay was benchmarked for RNA-based amplification using *A. minutum* as a representative species. To assess the limit of detection (LoD) and quantitative range, a synthetic RNA fragment corresponding to the target region of the *A. minutum* consensus sequence (produced by Genewiz, UK) was used as input for a dilution series spanning 6 orders of magnitude. The equivalent test was then performed using total RNA of *A. minutum* VGO0657. Field-relevant RNA inputs were tested over 4 orders of magnitude spanning 0.1 – 100 ng per reaction, without prior information on the *sxt*A4 transcript content of the cells.

For specificity testing we included four strains of *Alexandrium minutum* (FE126, CCAP 1119/48, VGO0657, VGO0650) as positive controls, another SXT producing dinoflagellate *Gymnodinium catenatum* (VGO1486), and four representative non-dinoflagellate phytoplankton species. For these non-dinoflagellate species, genomic DNA was tested directly. All tests were performed using inputs of 10 ng reaction^-1^.

### 2.8 Determination of *sxt*A4 transcripts per cell in *A. minutum* strains

We determined the number of *sxt*A4 transcripts per cell under nutrient replete condition in three cultured strains with different cell volume and growth rate profiles (strains VGO0657, CCAP 1119/48 and FE126). Growth rates were calculated by bi-daily counting of triplicate culture flasks in the middle of the light phase when cells were not dividing. The specific growth rate *μ* (cells day^-1^) during the exponential phase was calculated using the formula *μ* = (ln(N_1_)-ln(N_0_)) / (t_1_ - t_0_) (Anderson et al., 1990), where N denotes cell concentration and t denotes day of culture between two timepoints. Cells were then kept in exponential growth phase for at least 10 generations for stable physiology. Cells were collected for RNA extraction in mid-exponential phase using 5 μm pore size polycarbonate filters to exclude bacterial cells (Cytiva, US). Cell size of the 3 strains was also measured using an Operetta CLS High-Content Analysis System (Revvity).

From these samples we determined total RNA cell^-1^ as well as *sxt*A4 transcripts cell^-1^ using both our qRT-RPA assay as described above and qRT-PCR primers that we specifically designed for this purpose (Table 1). The qRT-PCR primers uSxtA4_3264F - uSxtA4_3410R, overlap with the RPA probe and reverse primer for comparability with the RPA assay, and amplify with 86% efficiency based on a standard curve analysis. The 20 μL reactions volumes contained 1x GoTaq qPCR Master Mix (Promega, UK), 1x GoScript™ RT Mix (Promega, UK), 200nM each primer and 2μL template. The cycle was: 1x (42°C for 15 minutes), 1x (95°C for 10 minutes), 40x (95°C for 30s, 64°C for 30s, and 72°C for 30s) on a CFX Opus 96 Real-Time qPCR system (BioRad, USA).

### 2.9 Assay performance with in simulated samples

Validation of the assay involved the production of simulated samples, as a bloom did not occur in our planned sampling area during the study period. We performed three tests: 1) Spiking target RNA into a mixture of non-target RNA from cultured species, 2) Spiking target RNA into RNA extracted from natural seawater, and c) spiking cultured cells into natural seawater and then extracting the total community RNA.

The first test was performed for three *A. minutum* strains (FE126, CCAP 1119/48, VGO0657). Spike RNA was added to 1 ng reaction^-1^ into a mixture of non-target RNA composed at 10 ng reaction^-1^. This background was made of 4 equal parts (2.5 ng reaction^-1^ each) of the green alga *Bathycoccus sp.,* the coccolithophore *Gephyrocapsa (Emiliania) huxleyi* and the diatoms *Phaeodactylum trichornutum* and *Thalassiossira pseudonana*.

In the second test, a dilution series of *A. minutum* CCAP 1119/48 total RNA was performed using RNA from a natural seawater sample. A 1.5 L water sample was collected from Karteros Beach, 10 km east of Heraklion, Crete, Greece on the 3^rd^ of July 2025, containing non-target dinoflagellate species including *Peridiniales*, *Gymnodinium*, *Tripos* spp., *Prorocentrales*, *Protoperidinium* and *Gonyaulax,* as well as other phytoplankton and microzooplankton. The water sample was filtered through a 3.0 μm pore size polycarbonate membrane (Cytiva, US), snap frozen in liquid nitrogen and stored at −80 C until the RNA extraction. Again, this background RNA matrix was kept at a fixed concentration of 10 ng reaction^-1^.

Finally, in the third test the assay’s performance was validated with spiking *A. minutum* VGO0657 cells in sea water sample. We collected 2 L seawater from Ammoudara beach in Heraklion, Crete, Greece on the 22^nd^ of November 2025. The sample was rich in diatoms and small naked dinoflagellates. Two mock samples were created by adding 1,000 and 10,000 cells into 0.4 L of the natural sample. Samples were filtered through a 3.0 μm pore size polycarbonate membrane (Cytiva, US) and RNA extraction, DNAse treatment and purification was performed as described for the cultured isolates. The total RNA was eluted in 10 μL water and 6.5 μL were added to the amplification reaction.

## 3 Results

In this study, we developed an RNA-based qRT-RPA assay to quantify expressed *sxt*A4 from *Alexandrium* species. The main steps of the assay development are shown in **Figure 1**. Following a series of optimization to achieve the most efficient amplification, the performance across four dinoflagellate species was evaluated using *sxt*A4 PCR amplicons. Specificity was tested with several non-target species representing different phytoplankton taxa. The remaining characterization focused on *Alexandrium minutum* RNA as a reference species that is relevant for HABs globally, and was physiologically stable in our culture conditions. Amplification range and sensitivity was determined using both total RNA, where *sxt*A4 copies per cell were unknown, as well as synthetic RNA to enable *sxt*A4 transcript quantification across different *A. minutum* strains. Analysis of three strains with differing size, total RNA content and *sxt*A4 transcript abundance provide an approximate relationship of *sxt*A4 transcripts to cell abundance, to demonstrate relevance to HAB thresholds. Finally, to demonstrate the field applicability of the assay, yet in the absence of a bloom during the study period, the assay performance was tested using simulated samples. These were prepared either by adding target RNA into a background RNA from natural samples, or adding target cells to natural seawater and treating it as a simulated field sample.

**Figure 1.**
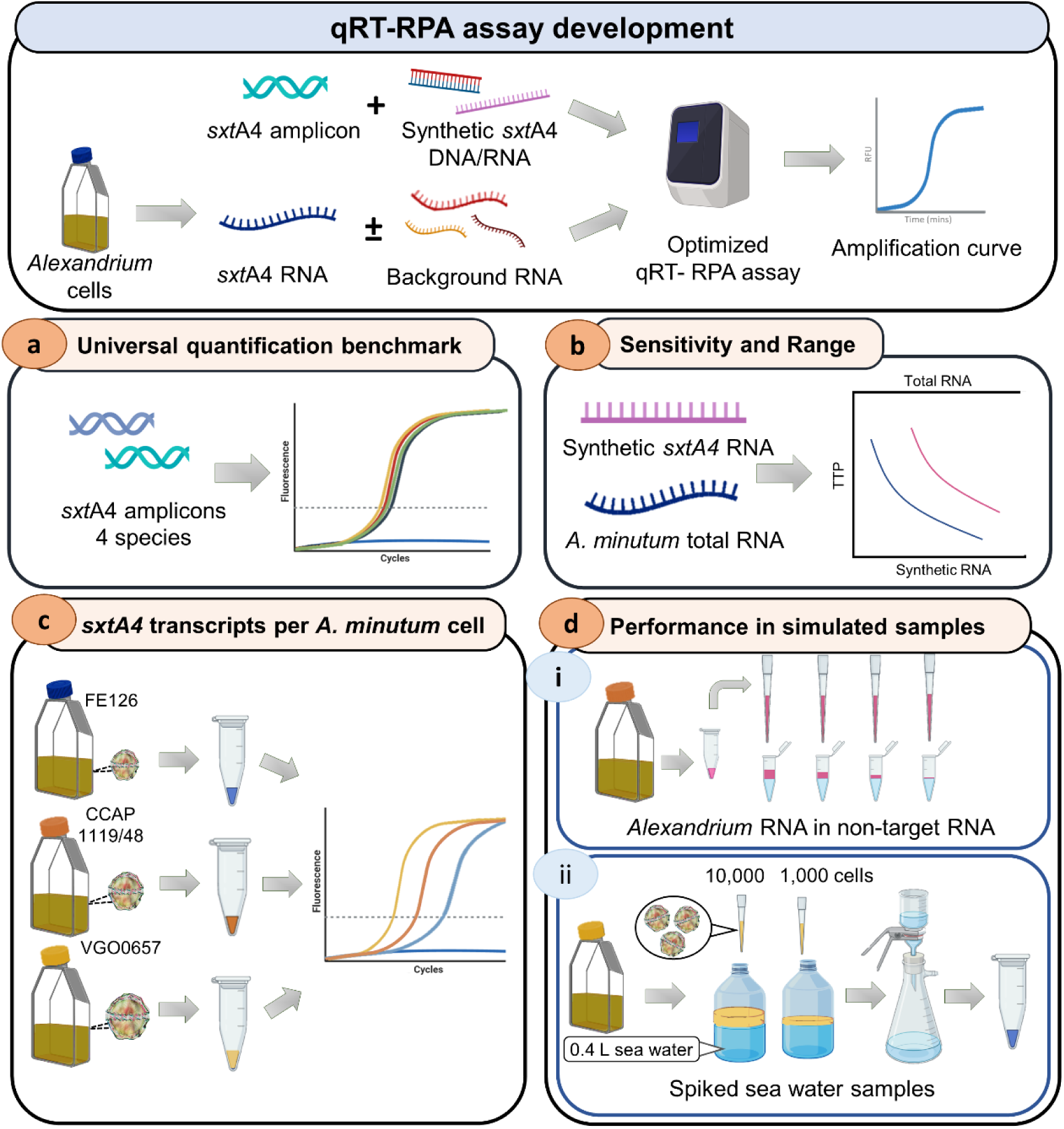
Key steps in qRT-RPA assay development. Assay optimization used amplicons and synthetic templates, with further benchmarking focusing on *A. minutum*. (a) Assay performance across *Alexandrium* species using *sxt*A4 amplicons. (b) Quantitative range of the assay using synthetic *sxt*A4 RNA and total *A. minutum* VGO0657 RNA. (c) Calculation of *sxt*A4 transcripts per cell in 3 different *A. minutum* strains using qRT-RPA assay. (d) Simulated samples including (i) spiking a dilution series of *A. minutum* CCAP 1119/48 RNA into background RNA and (ii) spiking two concentrations of *A. minutum* VGO0657 cells in a sea water sample. The image was created with BioRender.com.

### 3.1 Primer and probe design

The assay design was targeted to the first ∼ 400 bp part of the *sxt*A4 open reading frame which contains a highly conserved region with the least nucleotide polymorphisms among *Alexandrium* species. This is key for a truly “universal” RPA probe design, as even a single mismatch can reduce or preclude target amplification. Mismatches are better tolerated in the primers than in the probe (Liu et al., 2019). Nevertheless, we avoided regions with polymorphisms at the 3’ end of the primer or used degenerate bases to encompass possible sequence variation (**Figure 2**). We tested 2 forward primers, in combination with a single reverse and probe. By using amplicons of the *sxt*A4 region from four different species, we could compare amplification based solely on sequence variation, by conducting tests at a single template input concentration of 10^6^ copies per reaction.

**Figure 2.**
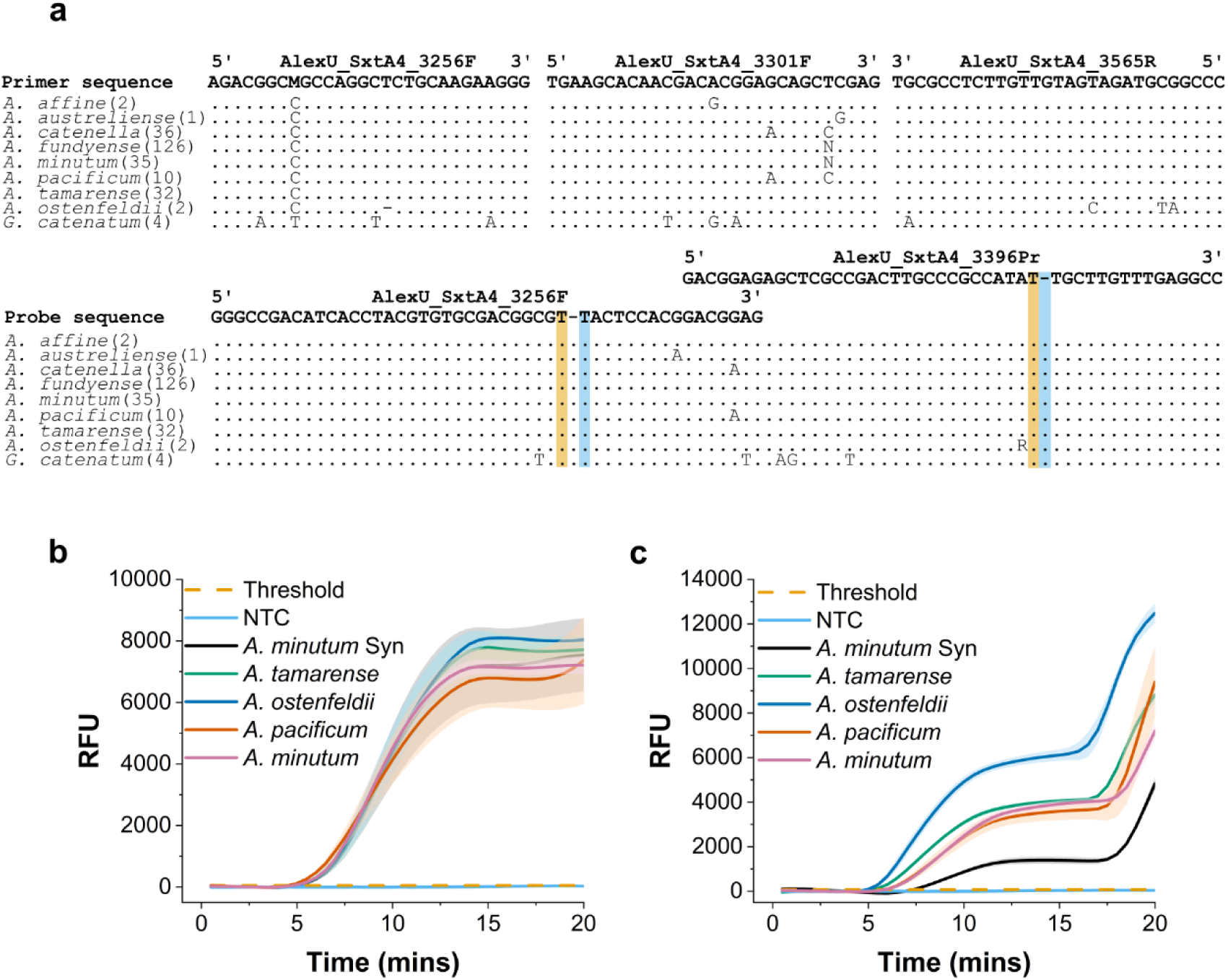
qRT-RPA assay design and testing. a) Nucleotide alignment of *Alexandrium* and *Gymnodinium catenatum sxt*A4 sequences from NCBI, along with the tested universal *sxt*A4 primers and probes. Numbers in brackets denote the number of sequences used to generate the consensus sequence for each species. Multiple alleles are represented as degenerate bases. The probe fluorophore and quencher positions are highlighted in orange and blue, respectively; the dash represents the abasic site. b) qRT-RPA amplification curves using and AlexU_SxtA4_3256F / AlexU_SxtA4_3565R / 3396Pr. c) qRT-RPA amplification curves using AlexU_SxtA4_3301F / AlexU_SxtA4_3565R / 3396Pr. The notation Syn denotes synthetic RNA generated based on the *A. minutum* sequence. All amplification tests were done using amplicons of the target regions at 10^6^ copies reaction^-1^.

Despite mismatches in the 3’ end of the forward primer AlexU_SxtA4_3301F, *sxt*A4 amplified from all *Alexandrium* species. However, TTP differed among species and those with no mismatches amplified faster. The primer and probe combination AlexU_SxtA4_3256F / 3565R / 3396Pr amplified a 309 bp region from all species with an equal TTP. This was selected as the Universal qRT-RPA assay and taken through further optimization.

### 3.2 qRT-RPA assay optimization

To maximise assay sensitivity and speed, we performed optimization tests for the reaction temperature, primer and probe concentrations, and MgOAc levels, based on the ranges of tolerance suggested by the manufacturer. These tests were performed using the synthetic DNA template representing *Alexandrium minutum sxt*A4, prior to inclusion of the reverse transcriptase needed for RNA amplification. The template input was invariable 10^4^ copies per reaction, which was near the initial limit of detection using RPA’s default recommended parameters.

We performed optimization tests for 5 parameters individually using a sequential approach. At each step we selected the optimal condition and used it in the next optimization step – the conditions were tested in the order they are presented in **Figure 3a-f**. The final optimized assay consisted of the highest recommended probe concentration for a maximum fluorescence signal, the lowest possible primer concentrations that did not compromise amplification signal to minimize primer-dimer formation, the default MgOAc concentration a temperature of 40 ℃ which is favorable for the reverse transcription enzyme.

**Figure 3.**
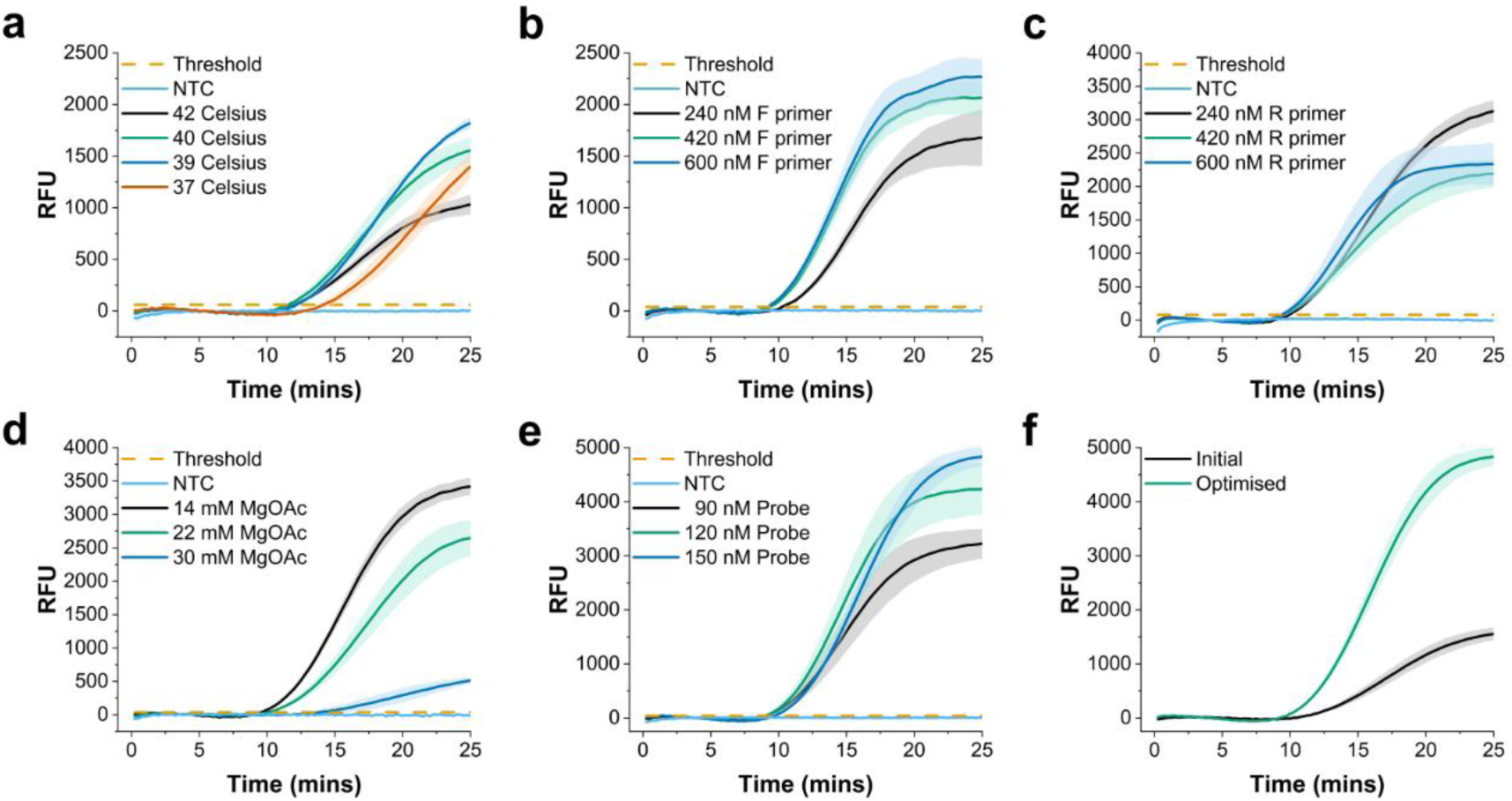
Assay optimization for different parameters using 10^4^ copies of *Alexandrium minutum* synthetic DNA template. (a) Temperature (b) Forward primer concentration (c) Reverse primer concentration (d) Magnesium acetate concentration (e) Probe concentration (f) Comparison between initial conditions based on manufacturer’s instructions and final optimized assay conditions. The coloured halo of each curve represents the SD from 3 technical replicates.

We selected the conditions yielding maximum signal intensity as the optimization strategy, rather than the quickest TTP. This accounts for the need to maintain a detectable signal at low concentrations for enhanced sensitivity. Also, it enables the assay to be run on portable devices that have less sensitive fluorescence detectors than laboratory-based qPCR cyclers. The final protocol for a 50 μL reaction was as follows: 29.5 μL rehydration buffer, 2.1 μL of 10 μM forward primer (420 nM final concentration), 1.2 μL of 10 μΜ reverse primer (240 nM final concentration), 0.75 μL 10 μM probe (150 nM final concentration) and 1 μL Superscript IV for RNA templates (Thermo Fisher Scientific, USA). These conditions resulted in the TTP becoming faster by 2.5 mins and a 3.3-fold increase in the maximum signal intensity (**Figure** 3**f**).

### 3.3 Amplification efficiency across multiple *Alexandrium* species

Assay optimization is normally initiated using gDNA as it is a more stable molecule and the type and amount of reverse transcriptase can be separately optimized. However, the *Alexandrium sxt*A4 does not amplify efficiently from gDNA, normally requiring addition of DMSO into PCR reaction. This not compatible with qRPA chemistry, making this approach unsuitable for *sxt*A4 quantification (**Figure S1**).

The performance of the optimized qRPA assay was evaluated using *sxt*A4 amplicons spanning a longer section of *sxt*A4 that contained the RPA assay target region. These were generated from four *Alexandrium* species: *A. tamarense* (RCC292)*, A. ostenfeldii* (CCAP 1119/53)*, A. pacificum* (CCAP 1119/52) and *A. minutum* (VGO0657) (**Figure 4**). This approach enabled direct comparison of amplification efficiency based solely on sequence variation, using accurately quantified inputs. Under optimized conditions, the assay showed uniform amplification kinetics across the four species and over four orders of magnitude, retaining quantitative performance in serial dilutions. The LoD was 10^3^ copies reaction^-1^ with the exception of the *A. tamarense* amplicon which failed to amplify below 10⁴ copies reaction^-1^. Overall, these results demonstrate robust, cross-species quantitative performance over a wide dynamic range.

**Figure 4.**
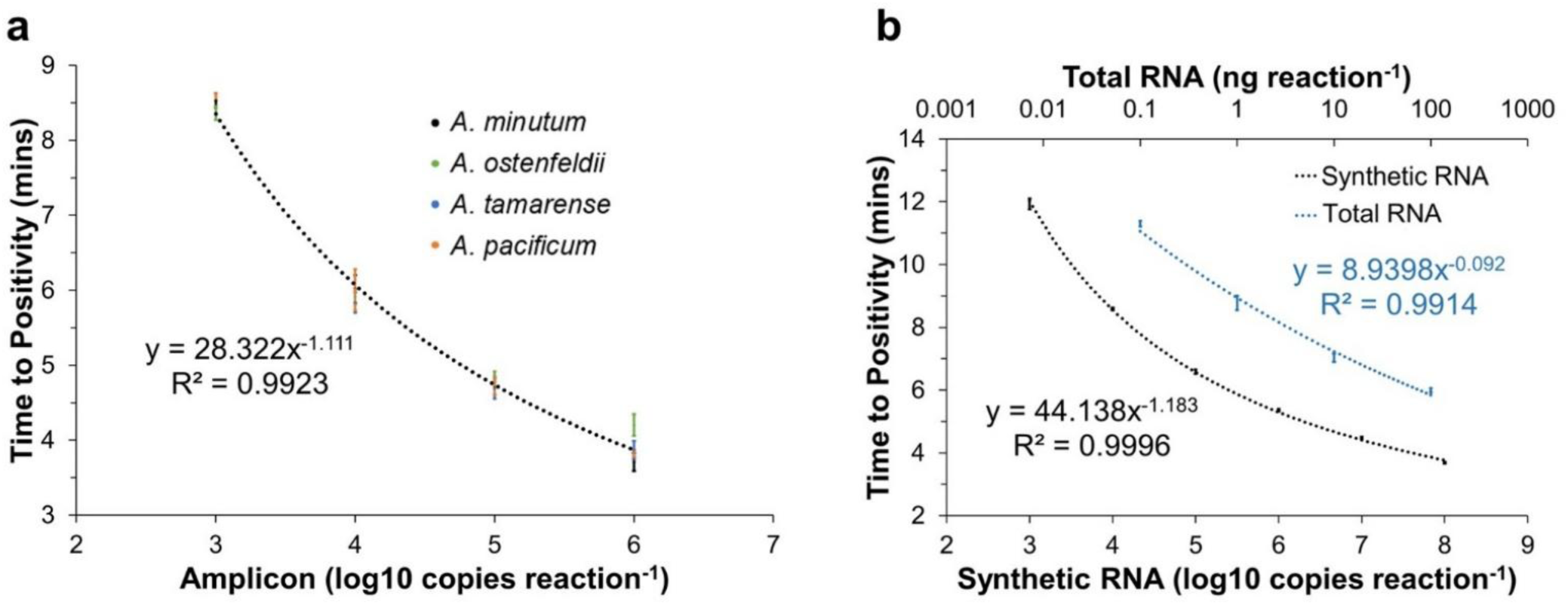
Characterization of the *sxt*A4 qRT-RPA universal assay. (a) Serial dilutions of isolated amplicons from four different *Alexandrium* species (*A. tamarense* (RCC292)*, A. ostenfeldii* (CCAP 1119/53)*, A. pacificum* (CCAP 1119/52)*, A. minutum* (VGO0657)). (b) Standard curves of *A. minutum* synthetic *sxt*A4 RNA (black) and *A. minutum* VGO0657 total RNA (blue). Error bars represent the standard deviation of 3 technical triplicates.

### 3.4 Sensitivity and quantitative range

Following the initial assay optimization using synthetic DNA and amplicon templates, the remaining assay characterization was performed on RNA templates using *Alexandrium minutum* as a reference strain. We assessed analytical sensitivity for RNA using both total RNA from *A. minutum* VGO0657 and synthetic *sxt*A4 RNA as a quantification standard (**Figure 4**). The assay’s quantification range spans at least 4 orders of magnitude and the limit of detection is 10^3^ synthetic RNA copies and 0.1 ng of *A. minutum* VGO0657 total RNA. Furthermore, the assay run time is fast, with the lowest concentrations of total RNA amplifying in under 12 minutes.

### 3.5 Assay specificity

We tested the RNA-based assay specificity using four strains of *A. minutum* as positive controls, the non-target saxitoxin-producing dinoflagellate *Gymnodinium catenatum* (VGO1486), and other non-target cosmopolitan microalgal species representing green algae, coccolithophores and diatoms (**Table 2**). Amplification in other *Alexandrium* species was already shown in previous tests using amplicons amplified from *sxt*A4 genomic DNA; active *sxt*A4 expression in these cultures was not tested due to unsatisfactory physiological state in our growth conditions. At the RNA level, the *sxt*A4 transcript was not detected in any non-target species.

**Table 2.** *Alexandrium sxt*A4 qRT-RPA assay specificity.

| Species | Strain ID | Phylum | *sxtA4* RNA |
| --- | --- | --- | --- |
| *Alexandrium minutum* | VGO0650 | Dinoflagellata | + |
|  | VGO0657 |  | + |
|  | CCAP119/48 |  | + |
|  | FE126 |  | + |
| *Gymnodinium catenatum* | VGO1486 | Dinoflagellata | - |
| *Bathycoccus sp.* | RCC4222 | Chlorophyta | - |
| *Emiliania huxleyi* | RCC6241 | Haptophyta | - |
| *Phaeodactylum tricornutum* | CCAP 1055/1 | Heterokontophyta | - |
| *Thalassiosira pseudonana* | CCMP 1355 | Heterokontophyta | - |

### 3.6 Relationship of *sxt*A4 transcripts to cell abundance

Trigger limits for HAB monitoring programs generally refer to specific cell concentrations. Therefore, we sought to equate our RNA-based assay results to equivalent cell number. For this purpose, we determined the number of *sxt*A4 transcripts per cell for three *A. minutum* strains VGO0657, CCAP 1119/48 and FE126 which had differing growth rates and cell size (**Table 3, Figure S2**). Cells were collected during the exponential growth phase and samples of 100,000 cells were collected by filtration for each strain. We measured cell size using and *Operetta CLS High-Content Analysis System*, total RNA cell^-1^ by scaling RNA yield to the cell input, and transcript number cell^-1^ by qRT-RPA for defined total RNA amounts with their originating cell numbers. Cell diameter and cell volume varied 1.8-fold and 3.4-fold, respectively. The total RNA content, transcripts pg^-1^ RNA and transcripts cell^-1^, all increased with increasing cell size. Considering the qRT-RPA assay LoD of 0.1 ng reaction^-1^ *A. minutum* VGO0657 total RNA, this equates to the detection of 6 cells reaction^-1^.

**Table 3.**
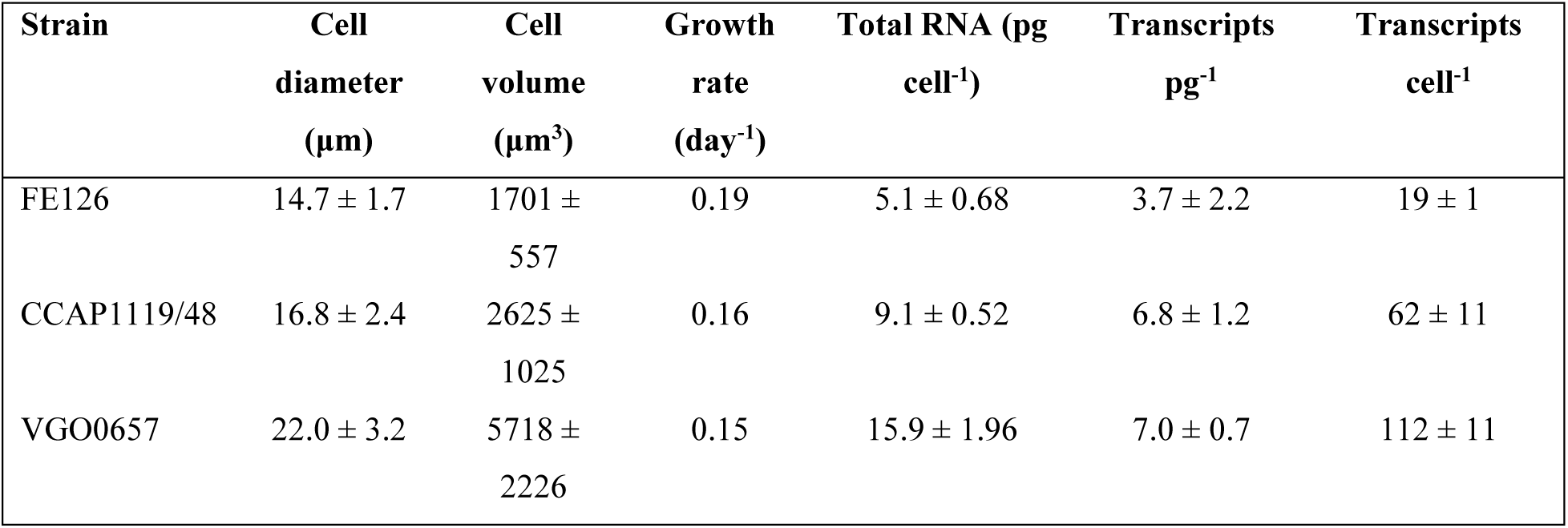
Cell size, total RNA content and *sxt*A4 transcript abundance analysis based on qRT-RPA for three *A. minutum* strains. Standard deviation is based on 3 replicates.

| Strain | Cell diameter (μm) | Cell volume (μm<sup>3</sup>) | Growth rate (day<sup>-1</sup>) | Total RNA (pg cell<sup>-1</sup>) | Transcripts pg<sup>-1</sup> | Transcripts cell<sup>-1</sup> |
| --- | --- | --- | --- | --- | --- | --- |
| FE126 | 14.7 ± 1.7 | 1701 ± 557 | 0.19 | 5.1 ± 0.68 | 3.7 ± 2.2 | 19 ± 1 |
| CCAP1119/48 | 16.8 ± 2.4 | 2625 ± 1025 | 0.16 | 9.1 ± 0.52 | 6.8 ± 1.2 | 62 ± 11 |
| VGO0657 | 22.0 ± 3.2 | 5718 ± 2226 | 0.15 | 15.9 ± 1.96 | 7.0 ± 0.7 | 112 ± 11 |

### 3.7 Assay performance in complex samples

We evaluated the influence of background RNA on *sxt*A4 RNA quantification by using sequentially more complex mock samples (**Figure 5**). Spiking 1 ng RNA from three *A. minutum* strains into 10 ng background RNA from a mixture of phytoplankton species (**Figure 5a**), showed a small degree of inhibition delaying the assay reaction by 1.7 minutes, on average, for the 3 strains. Based on the equations of VGO0567 synthetic and total RNA standard curves (**Figure 4b**), this would cause a quantitative underestimation of one order of magnitude.

**Figure 5.**
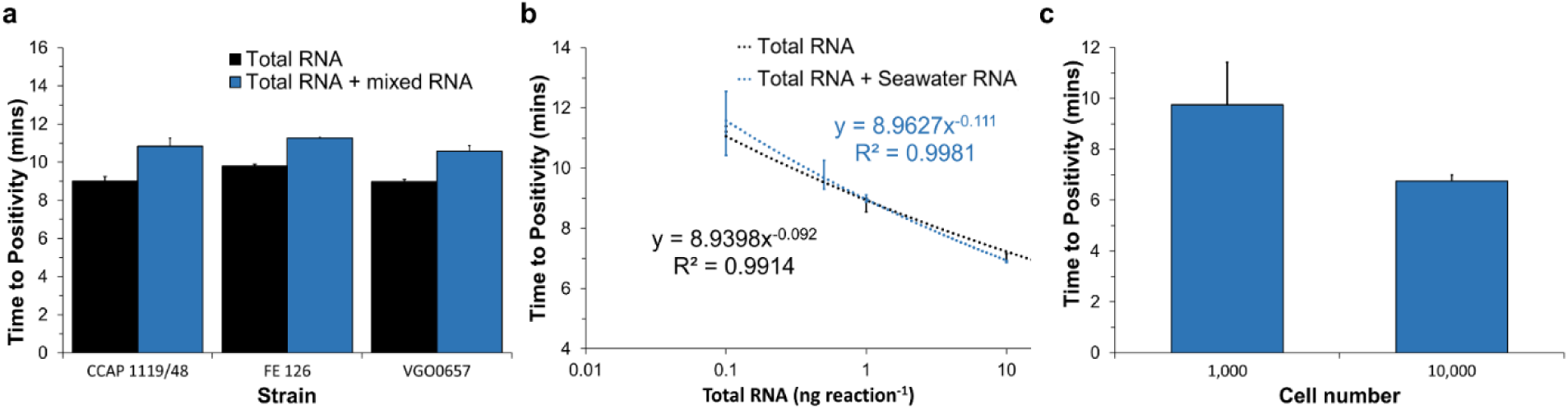
Assay performance in complex samples. (a) qRT-RPA of total RNA from *three A. minutum* strains (CCAP1119/48, FE126, VGO0657) in the presence (blue) and absence (black) of 10 ng background total RNA (b) qRT-RPA of total RNA from *A. minutum* CCAP 1119/48 in the presence (blue) and absence (black) of 10 ng total RNA isolated from a natural seawater sample. (c) qRT-RPA of total RNA from 1,000 and 10,000 cells of *A. minutum* VGO0657 spiked in sea water.

To further test this in more realistic conditions, we used 10 ng total RNA from a natural sample, spiked with *A. minutum* total RNA ranging 0.1 – 10 ng reaction^-1^. For this test we used *A. minutum* CCAP 1119/48, which showed healthier growth at the time of the experiment. Also, it is the representative strain with average size and *sxt*A4 transcript content. In this test, no inhibitory effect was observed near the assay’s LoD at 0.1 ng, however, amplification was more variable resulting in a higher standard deviation. The final LoD of the assay in complex samples was therefore set as 0.1 ng reaction^-1^ of *A. minutum* CCAP 1119/48 total RNA. According to the information in **Table 3**, this is equivalent to 11 *A. minutum* CCAP 1119/48 cells reaction^-1^. This LoD is indicative and expected to vary according to the strain *sxt*A4 transcript content, both intraspecifically and depending on physiology.

Finally, we tested the assay performance using mock seawater samples spiked with 1,000 and 10,000 cells of *A. minutum* VGO0657 (**Figure 5c**). Detection of 1,000 cells was easily achieved within the assay’s 12 min run time. The TTPs for samples spiked with 1,000 cells and 10,000 were 9.75 and 6.75 mins respectively, which based on the equation shown in **Figure 5b** (blue, with background RNA) equates to 0.5 ng and 12.9 ng total RNA per reaction respectively. This approximates one order of magnitude difference in the cell inputs.

## 4 Discussion

### 4.1 Methodological advance

This study demonstrates the utility of qRT-RPA for quantification of active toxigenic *Alexandrium* species. Targeting RNA enables preferential detection of cells that are actively expressing genes required for toxin production, while also improving analytical sensitivity due to the potentially higher abundance of transcripts relative to genomic copies (Geffroy et al., 2021; Murray et al., 2015). Amplification is achieved very fast in less than 12 mins, compared to approximately one hour for qPCR and ddPCR. Importantly, amplification at a constant low temperature around 40 °C, makes qRT-RPA ideal to use with simple portable devices. Furthermore, the tolerance of qRT-RPA to minor interspecific sequence variation, resulted in quantification of *sxt*A4 amplicons representing multiple *Alexandrium* species with equal efficiency. This ensures applicability across different geographic locations, where the causative microorganism might vary or where multiple saxitoxin producing species may coexist. The speed, ease of assay design and energy efficiency makes qRT-RPA ideal for RNA-based testing for toxigenic *Alexandrium* species in the field, enabling functional gene monitoring for early-warning HAB surveillance.

### 4.2 Performance characteristics

The qRT-RPA assay exhibited a quantitative range spanning four orders of magnitude, with limits of detection of approximately 10³ synthetic *sxt*A4 RNA copies reaction^-1^ and 0.1 ng total *Alexandrium minutum* RNA reaction^-1^, corresponding to approximately 10-20 cell equivalents depending on strain. These sensitivities are comparable to those reported for DNA-based qPCR assays targeting the *sxt* gene, with limits of detection ranging from 2.5 to 150 cell equivalents reaction^-1^ (Gao et al., 2015b; Han-Sol et al., 2024; Murray et al., 2019), or <10 copies reaction^-1^ for purified DNA standards (Penna et al., 2015; Savela et al., 2016).

Cross-species testing using *sxt*A4 amplicons revealed uniform amplification kinetics across the *Alexandrium* species tested, despite some differences in the sequence. These results indicate that the assay design successfully minimizes sequence-dependent bias. Furthermore, the assay was highly specific to the target genus, with no amplification in SXT containing *G. catenatum*, other phytoplankton species, or natural samples containing diverse phytoplankton collected from the coast of Crete.

By validating assay performance using isolated *sxt*A4 amplicons rather than genomic DNA, we were able to separate sequence-dependent amplification behavior from confounding effects related to genome size and variable gene copy number, which are known to complicate molecular ecological analyses of dinoflagellates (Ruvindy et al., 2023). This approach revealed that the probe specificity was a key determinant for equivalent performance of qRT- RPA across different species. This is consistent with previous studies on RPA probe mismatch sensitivity (Liu et al., 2019).

Background non-target nucleic acids are known to have an inhibitory effect on RPA, most likely due to non-specific primer binding at the low running temperature (Munawar, 2022; Rohrman and Richards-Kortum, 2015). In our assay, we found moderate inhibition in the presence of complex background RNA from cultured phytoplankton, which was expected as one of the primers is degenerate (Liu et al., 2019). The quantitative underestimation equated to less than one order of magnitude. However, this effect did not persist when a more complex natural sample was used as the background RNA. Perhaps a larger diversity of sequences causes a dilution of potentially interfering sequences. Our results highlight that, standard curves with natural background RNA are more representative for future quantification of the target in unknown samples, and should be adopted as a standard practice for qRT-RPA assays. Importantly, reliable detection at low target concentrations was achieved.

Although translating *sxt*A4 RNA-based detection to cell equivalents is desirable for comparability to established monitoring programme thresholds, this will vary considerably with strain and physiological state. Nevertheless, detection of as little as 11 cell equivalents reaction^-1^ from cells grown in optimal conditions, usually associated with low toxicity (Bui et al., 2024), suggests that the assay is likely to be highly sensitive for actively toxigenic field populations. Furthermore, this limit of detection is highly suitable for the target ranges of HAB monitoring programmes with the strictest alert thresholds from 100 *Alexandrium* cells L^-1^.

### 4.3 Comparison with other isothermal amplification approaches

Isothermal amplification strategies for HAB detection have been explored primarily aiming at field deployability. Most reported approaches rely on DNA-based detection with qualitative or endpoint readouts, including RPA or LAMP coupled to lateral flow dipsticks or CRISPR–Cas signal amplification for species identification rather than functional risk assessment (e.g. haptophytes, *Karenia mikimotoi*, *Pseudo-nitzschia multiseries* (Wang et al., 2023; Yao et al., 2024; Yu et al., 2025; Zhang et al., 2024). A notable exception is the use of RPA coupled to Nanopore sequencing, which enables portable, high-resolution taxonomic profiling but remains non-quantitative and DNA-based (Hatfield et al., 2020). Among isothermal methods, the only study to date targeting a toxin biosynthesis gene rather than a ribosomal marker is a LAMP assay targeting *dab*A in *Pseudo-nitzschia multistriata*, demonstrated on a custom-made field-deployable device; however, it is also DNA-based and lacks detailed quantitative and specificity characterization (Alrefaey et al., 2026). In contrast, the qRT-RPA assay presented here uniquely combines isothermal amplification, transcript-level detection and real-time quantification, making it well suited for early-warning applications where speed, functional relevance and portability are prioritized.

### 4.4 Conclusion

This study establishes the analytical performance of a universal qRT-RPA assay targeting the *sxt*A4 transcript, enabling rapid, cross-species detection of toxigenic *Alexandrium* with functional relevance and short time-to-result. Validation under controlled conditions, including sequence diversity and increasing matrix complexity, provides a solid methodological foundation for RNA-based isothermal monitoring. As with any emerging molecular approach, targeted field testing with natural bloom samples and direct comparison to qPCR, microscopy and toxin measurements will be required to define operational performance and establish informed risk thresholds. With full compatibility to low-power portable fluorescence readers and sensor-integrated platforms, qRT-RPA is well positioned as a rapid early-warning tool in HAB monitoring programmes.

## Acknowledgements

The authors would like to thanks Dr. Nuria Lluch, Dr. Rosa Isabel Figueroa and Dr. Esther Garces at Insitut de Ciencias del Mar, Spain, for contributing *Alexandrium minutum* and *Gymnodinium catenatum* cultures. This research was funded by the EU Horizon project AquaBioSens no. 101135432.

