## Supplementary Information for "Universal rapid RNA-based quantification of toxigenic *Alexandrium* species (Dinophyceae) using quantitative recombinase polymerase amplification"

### Supplementary Figures

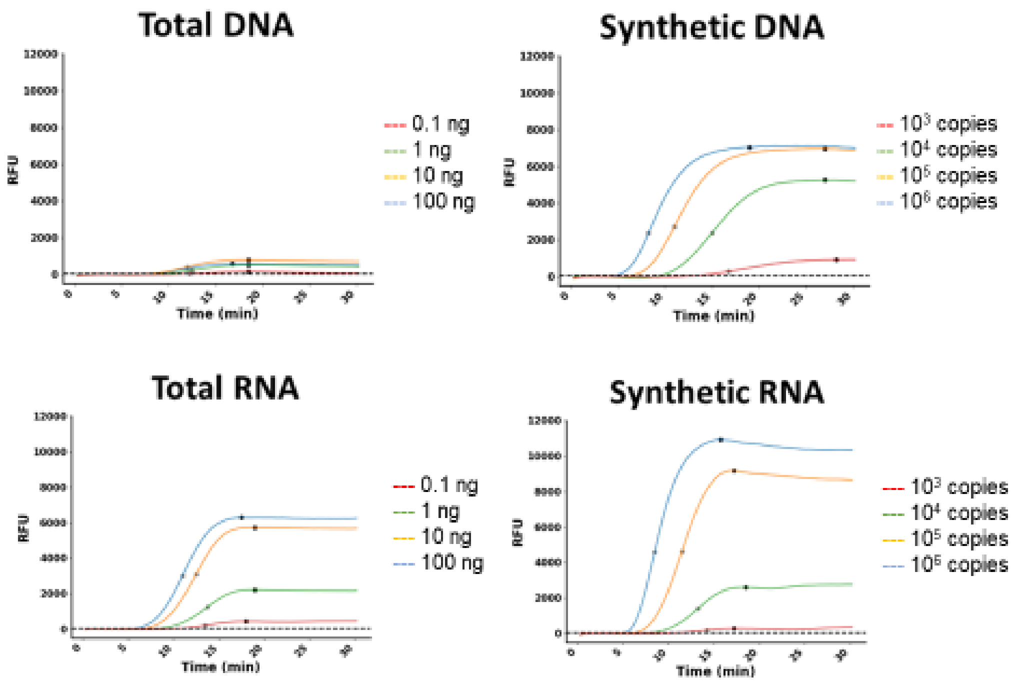


Figure S1. Performance of the universal *sxt*A4 assay with different *A. minutum* templates over a range of inputs concentrations per reaction. Total (genomic) DNA show less efficient amplification relative to other template types.

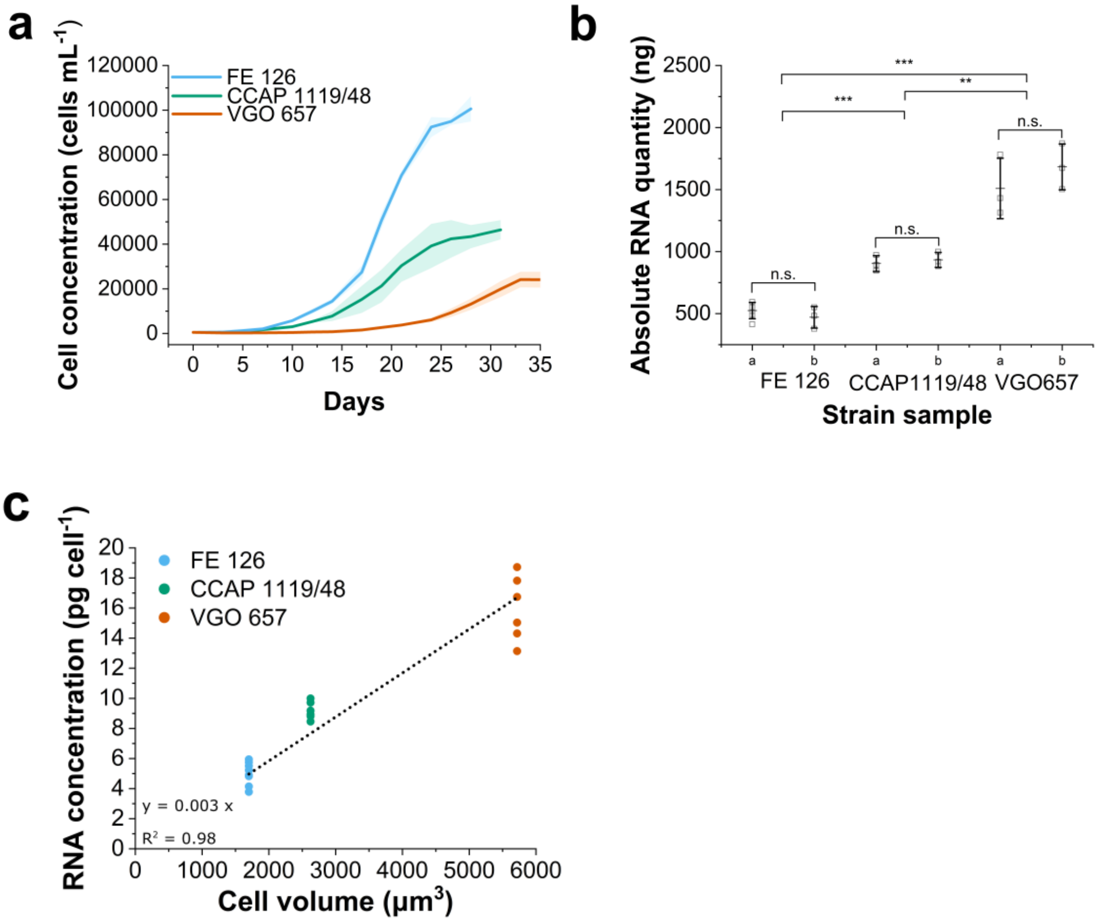


**Figure S2**. Total RNA yield and cell size analysis of *A. minutum* strains FE126, CCAP1119/48, VGO657. (a) *sxt*A4 qRT-RPA amplification curves. (b) Yield from RNA extraction of 100,000 cells. Samples a and b from each strain were both collected and extracted on different days. (c) Correlation of the average cell volume with RNA concentration per cell calculated from b. * = $P<0.05$, ** = $P<0.01$, *** = $P< 0.001$
